# Integrated multi-omic and functional evidence uncover substantial strain-specific divergence in zebrafish

**DOI:** 10.1101/2025.10.24.684330

**Authors:** Wenjun Chen, Jing Wang, Ziyi Li, Jianbo Jian, Yang Liu, Lingling Zheng, Shunping He, Liandong Yang

## Abstract

The zebrafish (*Danio rerio*) is a foundational vertebrate model in biomedical and developmental research, yet genetic divergence among laboratory strains remains an underappreciated source of biological and experimental variation. Among these, the AB and Tübingen (TU) strains are used globally, forming the basis for most zebrafish-based discoveries. Here, we integrated long- and short-read sequencing, ATAC-seq, Hi-C, transcriptomics, and functional assays to generate a comprehensive multi-omic comparison of these two strains. Our analyses revealed substantial genomic and regulatory divergence, including a 62-bp insertion in the *asmt* promoter that elevates transcription and alters melatonin biosynthesis, driving strain-specific circadian phenotypes. We also identify a non-synonymous variant (*mybbp1a*^G2612C^) associated with heightened immune responsiveness. Together, these findings construct the first genome-wide atlas of functional variation between zebrafish strains, illuminating how hidden genetic differences can shape phenotype, regulatory architecture, and experimental reproducibility in this essential vertebrate model.

## Introduction

*Danio rerio* (zebrafish) has become a widely used model organism in genomics^1^, behavior^2^, immunology^3^, toxicology^4^, neuroscience^5^, developmental biology^6^ and human diseases^7^. Its popularity stems from several advantages^1,8^, including a high reproductive rate, short life cycle, transparent embryos for easy observation, suitability for genetic manipulation, compatibility with various research techniques, and significant genomic similarity to humans. Inbred zebrafish strains, such as AB, TU, WIK, are used in over 1,200 laboratories worldwide^9,10^. These strains have distinct genetic backgrounds, leading to notable genetic variation and phenotypic differences, ^11–15^ including variations in circadian rhythm behavior^16^, susceptibility to intestinal inflammation^17^, learning and memory^18^, and growth performance^19^.

Genetic variation is typically classified into three main categories based on the number of nucleotides alterations: single-nucleotide polymorphisms (SNPs), short insertions or deletions (InDels; 1–50 bp), and structural variants (SVs; ≥50 bp), including insertions (INS), deletions (DEL), inversions (INV), and duplications (DUP)^20,21^. Genetic variations influence phenotypes by altering protein structure, regulating gene expression, or disrupting RNA splicing^22–26^. Compared to SNPs and InDels, SVs may induce more significant phenotypic changes, as they can encompass multiple functional genetic elements^27^, directly alter gene content^28^, or affect the transcription levels of neighboring genes^29^. Additionally, SVs can directly regulate gene expression levels by altering gene copy number^30,31^. Given the pivotal role of genetic variation in phenotype formation, numerous studies have explored its impact on adaptive evolution, domestication, breeding, and disease. Specific research has focused on adaptive variation in high-altitude, hypoxic, and cold environments^32–35^, domestication of livestock^36–39^, breeding for desirable traits^29,40,41^, and the identification of variants linked to disease susceptibility^42–44^.

Over the past two decades, advances in genomics and sequencing technologies, along with the introduction of new computational algorithms, have greatly enhanced the discovery of genetic variation. Early methods, such as chromosomal microarrays (CMA) for detecting copy-number variations (CNVs)^45^, were followed by the use of short-read sequencing (SRS) technology to detect SNPs and InDels detection^46^, More recently, long-read sequencing (LRS) technologies have been developed and applied for the phasing and analysis of complex SVs^47^. Among these, LRS technologies, including PacBio single-molecule real-time (SMRT) sequencing and Oxford Nanopore Technologies (ONT) sequencing, have become widely used for SV detection due to their long, contiguous reads that can span SV breakponits^36,42–44,47^.

Through the systematic integration of LRS and SRS technologies, we comprehensively mapped the genetic variation landscape across the zebrafish AB and TU strains, including SNPs, InDels and SVs. Population genomic analyses revealed significant genetic divergence between AB and TU, consistent with distinct subpopulation structures. Our results support the following model: variation between strains is closely linked to epigenetic inheritance. Specifically, (1) differential accessible chromatin regions (dACRs) are more likely to be associated with highly differentiated SNPs; (2) SVs tend to perturb TAD structures, reshaping regulatory relationships and influencing gene expression. Our analyses also revealed highly concordant selection signatures across SNPs, InDels, and SVs, with top outlier variants frequently converging on the same genomic regions and nearby genes. These findings suggest that genetic variation at multiple scales collectively contributes to inter-strain divergence. By integrating multi-omics datasets—including behavioral assays, physiological measurements, RNA-sequencing and immune profiling—with whole-genome variant annotation, we identified a 62 bp insertion upstream of *asmt* and a non-synonymous SNP in *mybbp1a* as potential causal drivers of strain-specific phenotypic variation. These results establish a functional link between polymorphic architectures and adaptive traits, enhancing our understanding of how combinatorial genetic variation shapes population-specific phenotypes in vertebrate models.

## Results

### Identification and characterization of genetic variants in AB and TU zebrafish strains

LRS data were generated from 22 TU strain and 22 AB strain individuals using nanopore sequencing, with an average coverage depth of 24× (Supplementary Table 1 and Supplementary Figure 1). SVs were detected for each genome relative to the GRCz11 reference genome using multiple SV callers specifically designed for LRS data (Supplementary Table S2). On average, 77,475 SVs were detected per sample (Figure 1a). After filtering out unreliable genotypes based on a hard threshold, a total of 178,894 unique SVs were identified, including 222 DUPs, 103 INVs, 76,113 DELs, and 102,456 INSs (Figure 1c). The growth of the non-redundant SV set was initially steep but plateaued as additional samples were added. Similarly, the number of shared SVs decreases rapidly initially and then stabilizes as additional samples are included, suggesting that this subset of 44 zebrafish captures a large proportion of the common SVs (Figure 1b). All subsequent analyses were performed based on this SVs set.

**Figure 1.**
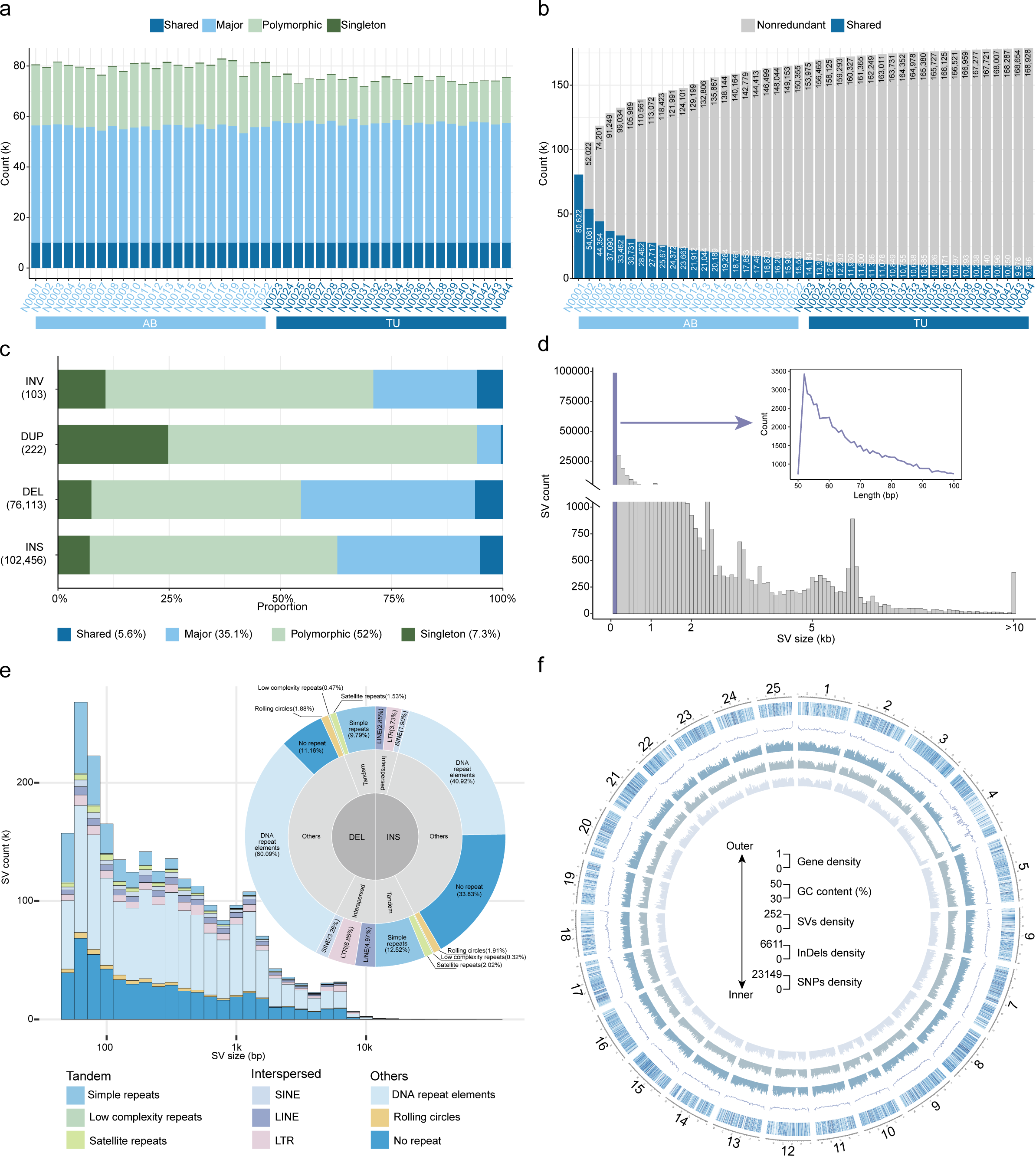
SVs discovery in 44 AB and TU strain zebrafish. **a.** The number of SVs for each discovery category is shown per sample, including shared (identified in all samples, n = 44), major (present in ≥ 50% of samples, n = 22 to 43), Polymorphic (present in > 1 sample, n = 2 to 21), and singleton (identified in only one sample) SVs. **b.** SVs discovered from each sample were merged into a non-redundant set. Shared SVs are shown as purple in each bar. Light blue represents AB strain zebrafish, and dark blue represents TU strain zebrafish. **c.** The frequencies for each SV type: inversion (INV), duplication (DUP), insertion (INS), and deletion (DEL). **d.** Size of SVs identified in AB and TU strain zebrafish populations **e.** Distribution of insertions and deletions classified by intersected repeat elements. **f.** Distribution of the SNPs, InDels and SVs across whole genome.

The LRS-based SVs were classified into four categories: shared (identified in all samples, n = 44), major (present in ≥ 50% of samples, n = 22 to 43), polymorphic (present in > 1 sample, n = 2 to 21), and singleton (identified in only one sample). Our results show that polymorphic SVs constitute the vast majority, accounting for approximately 52% (Figure 1c). Additionally, the length distribution of SVs primarily falls within the 50 – 100 bp range (Figure 1d). We found that the breakpoints of SVs predominantly overlap with repetitive sequences, primarily DNA repeat elements and simple repeats (Figure 1e). This observation suggests that repetitive sequences may play a significant role in the formation of structural variations across the genome in zebrafish. Across the genome, SNPs, InDels, and SVs exhibit a co-distributed pattern. A pronounced depletion in centromeric regions was observed (Fig. 1f), consistent with lower accessibility and residual assembly gaps in centromeric DNA, which can reduce variant detectability^48^.

Using the Illumina platform, we sequenced samples from 22 AB and 22 TU strain zebrafish, achieving an average sequencing depth of 31× (Supplementary Table S3 and Supplementary Figure 1), which ensured the reliable detection of low-frequency variants, including SNPs and InDels. Based on SRS data, we identified a total of 39,580,820 SNPs across the 44 samples, including 551,719 in exons, 17,294,484 in introns, and 11,228,618 in intergenic regions, (Supplementary Table S4). Additionally, we quantified the number of InDels, identifying a total of 10,614,893 InDels, of which 32,030 were located in exonic regions. These 32,030 InDels were further classified into the following categories: 9,948 frameshift deletions, 8,303 frameshift insertions, 5,312 non-frameshift deletions, 4,324 non-frameshift insertions, 1,030 stopgain, 59 stoploss, and 3,054 unknown types (Supplementary Table S5).

### Genetic divergence and diversity between AB and TU Strains

We further investigated the genetic divergence and diversity between the AB and TU strains. To better characterize the genetic relationships of the zebrafish samples, we employed three complementary approaches using the SV dataset: construction of a neighbor-joining (NJ) tree, ancestry analysis with ADMIXTURE, and principal component analysis (PCA). In the NJ tree (Figure 2a), all 44 zebrafish individuals clustered into two distinct groups corresponding to the AB and TU strains. This population subdivision was further supported by the PCA (Figure 2b), which clearly separated AB and TU along the first principal component, and by ADMIXTURE analysis (Figure 2c), which assigned individuals into two ancestry components matching the strain identity. Consistently, population structure analysis based on the SNP dataset also revealed the same pattern (Supplementary Figure 2), with the AB and TU strains forming two well-defined genetic groups.

**Figure 2.**
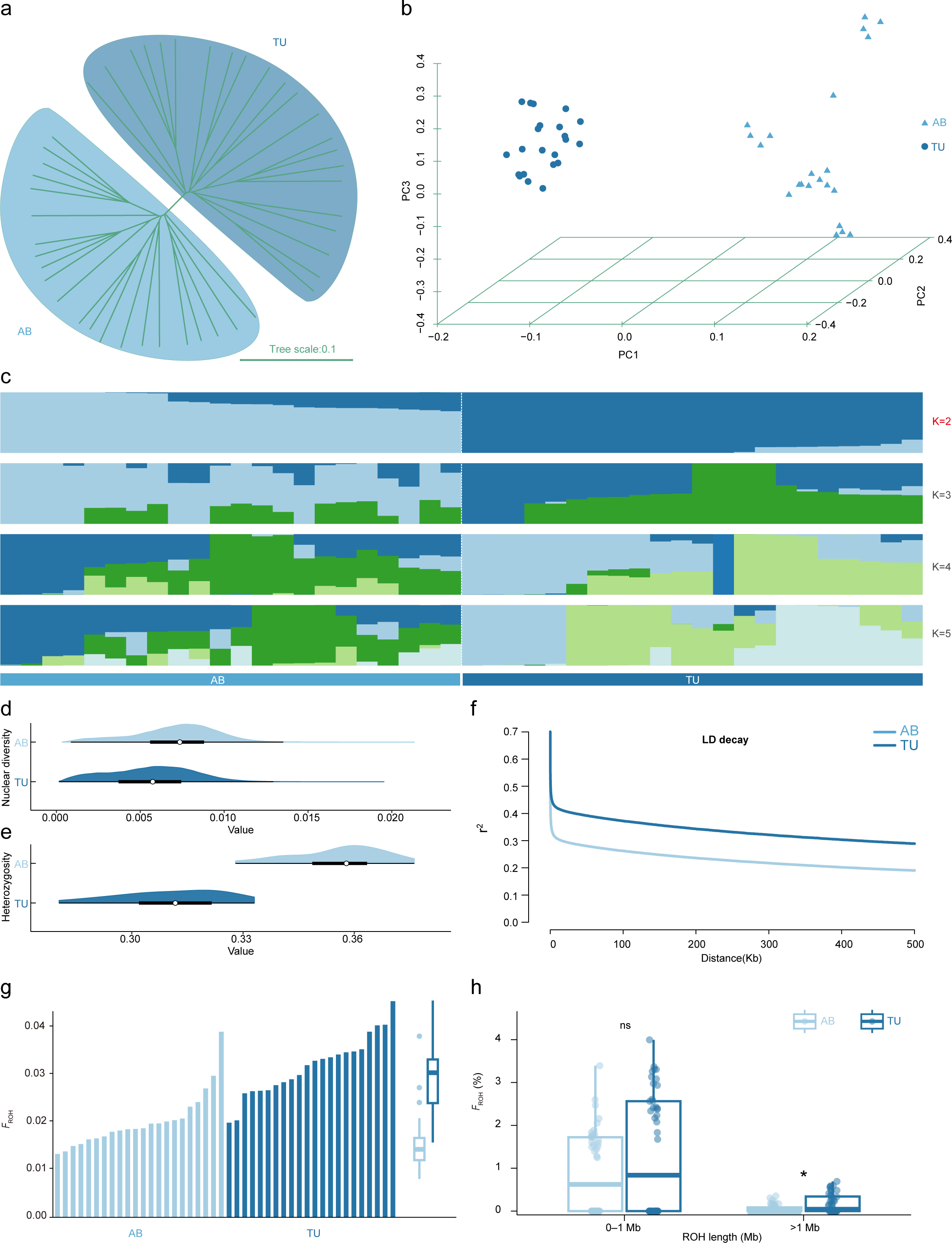
Phylogeny, population structure analysis and summary of the evaluation of genetic diversity of zebrafish strains AB and TU. **a.** Neighbour-joining tree based on genome wide SVs. Light blue branches indicate AB strain zebrafish and dark blue indicate TU strain zebrafish. The scale bar represents the genetic distance between individuals. **b.** Principal component analysis of 44 zebrafish using whole-genome SVs data. **c.** Genetic structure of AB strain zebrafish and TU strain zebrafish using the Admixture program based on SVs data. Each accession is represented by a bar, and the length of each coloured segment in the bar represents the proportion contributed by that ancestral population. When K=2, the cross-validation error (CV) value is the smallest. **d.** Comparison of nucleotide diversity (π) between two zebrafish strains. **e.** Mean individual genome-wide heterozygosity. **f.** Decay of linkage disequilibrium (LD) of AB strain and TU strain populations measured by the average LD coefficient r^2^. **g.** The distributions of inbreeding coefficient (*F*_ROH_). **h.** Comparison of *F*_ROH_ (%) across different ROH length categories between the AB and TU strains. The two-sided student’s t-test was applied and results were presented as mean ± SEM). *p < 0.05

To assess genetic diversity and demographic patterns in the two strains, we compared nucleotide diversity (π), heterozygosity, linkage disequilibrium (LD) decay, and runs of homozygosity (ROH) between AB and TU. Nucleotide diversity was lower in TU than in AB (Figure 2d), and a similar pattern was observed for heterozygosity (Figure 2e), indicating reduced within-strain genetic variation. Consistently, TU exhibited a slower decay of LD than AB (Figure 2f), reflecting a higher extent of haplotype homozygosity. Analysis of ROH showed that TU individuals harbor a significantly larger fraction of their genomes within homozygous tracts than AB (Figure 2f). When ROH were stratified by length, the difference was driven by long ROH (>1 Mb), for which TU exhibited higher *F*_ROH_, whereas no significant difference was detected for short ROH (<1 Mb). Combined with lower nucleotide diversity, reduced heterozygosity, and slower LD decay in TU, these patterns indicate reduced genetic diversity and elevated long-range homozygosity in TU, consistent with a smaller effective population size, potentially reflecting stronger inbreeding or historical bottlenecks during its breeding history.

### The Impact of Strain-Specific SNPs and SVs on Chromatin Accessibility and TAD Structure in Zebrafish

To investigate differences in chromatin accessibility between strains, we generated ATAC-seq data from juvenile fish of the AB and TU strains, each with two biological replicates. Replicates showed strong concordance around transcription start sites (TSSs) and clustered together in heatmap analyses (Supplementary Figure 3), which demonstrated the reproducibility of our data. Therefore, replicates within each strain were merged for downstream analyses. We identified 10,341–61,301 accessible chromatin regions (ACRs) per sample, covering 0.34%–2.90% of the GRCz11 reference genome (Fig. 3a). ACRs were classified into three categories based on their intersection with protein-coding genes: genic ACRs (overlapping gene bodies by ≥1 bp, 66.45%–69.43%), proximal ACRs (overlapping the ±2-kb flanking regions of genes by ≥1 bp, 2.13%–2.52%), and distal ACRs (all others, 28.30%–31.03%; Fig. 3b). We further partitioned ACRs into strain-specific and strain-shared sets (Fig. 3c).

**Figure 3.**
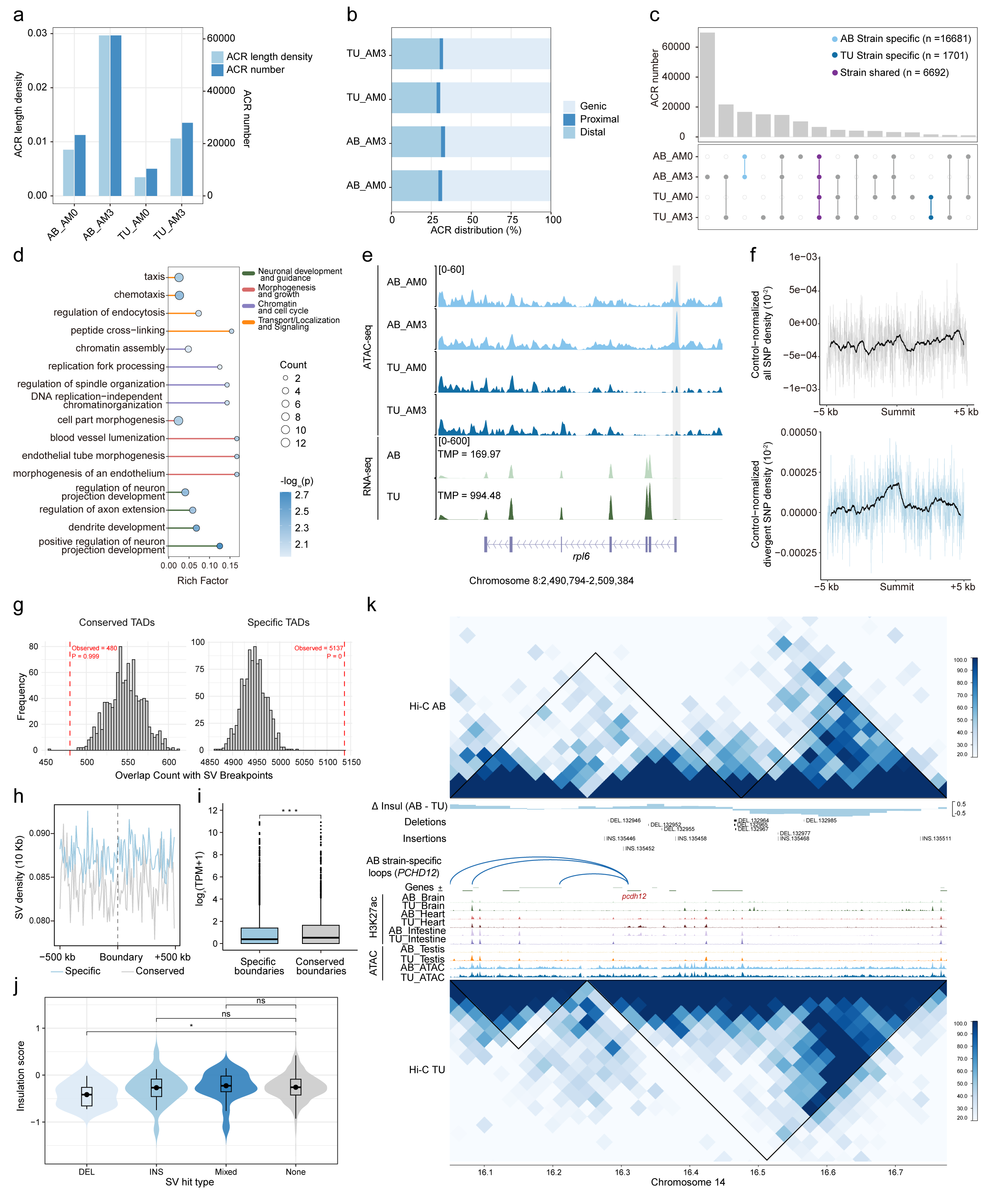
The Impact of Strain-Specific SNPs and Structural Variants on Chromatin Accessibility and TAD Structure in Zebrafish. **a.** The number and length of ACRs identified in each sample. **b.** The distribution of ACRs on different genomic features (genic, proximal and distal). **c.** Upset plot showing the ACRs that conserved or specifically existed across all samples. **d.** Gene set enrichment analysis of the nearest protein-coding genes to each dACR. The rich factor and -log_DD_(p) values are shown for various GO terms. Each point represents a GO term, with the size of the dot indicating the number of genes associated with the term. **e.** An example of dACR and RNA-seq data for the *rpl6* gene. **f.** SNP density around dACRs. The upper panel shows the control-normalized SNP density of all SNPs across a ±5 kb region around the dACR summit. The lower panel shows the control-normalized SNP density for divergent SNPs. **g.** Enrichment of SV breakpoints within conserved TADs and specific TADs. The distribution of overlap counts between SV breakpoints and TADs was computed based on 1000 random permutations (n = 1000 trials). The observed overlap count is shown by the dotted red line. The histograms represent the frequency distribution of overlap counts in both conserved and specific TADs, highlighting significant enrichment of SV breakpoints within. **h.** Density of structural variations around TAD boundaries. **i.** Expression of conserved and specific TAD boundaries. **j.** Insulation scores at TAD boundaries in relation to different SV hit types. The plot shows the insulation scores for deletion-type (DEL), insertion-type (INS), mixed-type (Mixed), and unhit (None) SVs. **k.** A potential example of SVs disrupting TAD structure and their association with AB strain-specific loops. The top panel shows Hi-C data for the AB strain, with insulation scores (Δ Insul) indicating differences between AB and TU strains. The middle panel displays SVs (deletions and insertions) in the region, with AB strain-specific loops highlighted around the *pcdh12* gene, which is differentially expressed in two strains. H3K27ac ChIP-seq and ATAC-seq tracks are shown across the region. The bottom panel shows Hi-C data for the TU strain, where the AB strain-specific loops are not within the same TAD. The two-sided student’s t-test was applied and results were presented as mean ± SEM). *p < 0.05, **p < 0.01 and ***p < 0.001.

We identified 477 strain-specific accessible chromatin regions (dACRs) between the AB and TU strains (Supplementary Table 6), defining the nearest protein-coding gene to each dACR as its associated gene. Gene set enrichment analysis revealed that these genes are primarily involved in processes such as neuronal development and guidance as well as morphogenesis and growth (Figure 3d) (Supplementary Table 7). By integrating dACR associations with transcriptomic data, we highlighted *rpl6*, a gene known to be involved in zebrafish brain development^49^, as a candidate. Notably, a dACR overlaps the *rpl6* promoter, and differential expression of *rpl6* between the two strains suggests that differences in promoter accessibility may contribute to its transcriptional divergence (Figure 3e). We further investigated whether SNPs are associated with dACR formation. First, we found no enrichment of all SNPs at dACRs, as shown by the uniform distribution of SNP density around the dACR summit (Figure 3e upper panel). However, we observed significant enrichment of divergent SNPs in dACRs (Figure 3e lower panel), suggesting a potential link between strain-specific SNP divergence and the formation of dACRs.

Next, we explored the relationship between genetic variation and epigenetic regulation at a larger scale by integrating SVs dataset with previously published Hi-C data from AB and TU strains. We found that SVs are more likely to occur in strain-specific TADs rather than in conserved TADs (Figure 3g). Additionally, we observed that the SV density around strain-specific TAD boundaries is higher compared to conserved TAD boundaries (Figure 3h). We also found significant gene expression differences between genes located near conserved versus non-conserved TAD boundaries (Figure 3i). Notably, deletion-type SVs at TAD boundaries were associated with a significantly lower Insulation score compared to unhit boundaries (Figure 3j), suggesting that SVs may influence gene expression by altering TAD boundary insulation. We identified a potential example where SVs might disrupt TAD structure. In this region, we observe significant SV enrichment, and specifically, we found that AB strain-specific loops are associated with the differentially expressed gene *pcdh12*, which was reported associated with neurological diseases^50^. In the AB strain, *pcdh12* and its associated loops are located within the same TAD. However, in the TU strain, these loops are not within the same TAD, suggesting that SVs may have disturbed the TAD structure (Figure 3k). This disturbance could result in the formation of strain-specific regulatory relationships, leading to the differential expression of *pcdh12* between the two strains.

Using population-scale SRS and LRS data, we jointly profiled SNPs, InDels, and SVs to test whether different variant classes mark the same divergence hotspots between AB and TU. The composite Manhattan plot (Supplementary Figure 4a) showed numerous concordant peaks across the genome for all three classes, indicating co-localized signatures of differentiation. To quantify SNP divergence near SVs, we calculated the mean pairwise genome-wide *F*st value of surrounding SNPs in non-overlapping 5-kb windows centered on each SV and compared the top 1% SVs by *F*st-SV with the full SV set; flanking *F*st-SNP was significantly higher for the top-1% *F*st-SVs (Supplementary Figure 4b). Intersecting the top-1‰ *F*st windows from SNPs, InDels, and SVs identified for shared genes—*asmt* (Acetylserotonin O-methyltransferase), *ergic1* (Endoplasmic reticulum-Golgi intermediate compartment protein 1), *flt4* (Fms-related receptor tyrosine kinase 4), and *akap17a* (A-kinase anchoring protein 17A) (Supplementary Figure 4c)—highlighting candidate loci where multi-scale genetic variation converges during strain divergence.

### Positive selection signals between zebrafish strains

To identify positive selection signals between the AB and TU strains, we performed a comparative analysis using the SNP dataset, assessing *F*st, differences in nucleotide diversity (π AB/TU or TU/AB), and the cross-population composite likelihood ratio test (XP-CLR) within 10-kb sliding windows along the genome (Supplementary Figure 5). The top 5% of ranked windows common to all three independent methods were considered positive selection windows, with genes intersecting these windows classified as positively selected genes (PSGs). We identified 128 PSGs from 140 positively selected windows in the AB strain and 179 PSGs from 399 positively selected windows in the TU strain (Supplementary Table 8). Gene ontology (GO) analysis of the combined 307 PSGs revealed enriched categories related to immune and neural functions, including “regulation of T cell proliferation (GO:0042129)”, “regulation of mononuclear cell proliferation (GO:0032944)”, “regulation of leukocyte proliferation (GO:0070663)” and “regulation of presynapse organization (GO:0099174)” (Supplementary Table 9). Notably, several PSGs associated with circadian rhythm were identified, including *asmt*, *clock1a* (clock circadian regulator a), and *tph1a* (tryptophan hydroxylase 1 a), suggesting that variations in circadian rhythm-related pathways may also contribute to the phenotypic divergence between the two strains.

Using the SVs dataset, we identified 820 PSGs (Supplementary Table 10). Similar to the PSGs identified from the SNP dataset, these genes were predominantly enriched in functional categories such as “negative regulation of immune system process (GO:0002683)”, “regulation of CD4-positive, alpha-beta T cell activation (GO:2000514)” and “CD4-positive, alpha-beta T cell activation (GO:0035710)” (Supplementary Table 11). Several circadian rhythm-related genes exhibited signals of positive selection, including *asmt*, *clock1a*, *opn4xb* (opsin 4xb), and *opn5* (opsin 5). Additionally, we also identified PSGs associated with associative learning and memory, such as *cpeb3* (cytoplasmic polyadenylation element binding protein 3), *glra2* (glycine receptor, alpha 2) and *gng8* (guanine nucleotide binding protein, gamma 8), pointing to potential selection pressures on neural processes related to cognition and behavior.

### Effect of an upstream insertion in the *asmt* gene on zebrafish circadian behavior

Based on the identified selection signals, we yielded several PSGs associated with circadian rhythm. Previous studies have reported behavioral differences in circadian rhythm between the two zebrafish strains^16^. To further investigate the circadian rhythm difference between the AB and TU strains, we conducted behavioral experiments. Our results showed that the AB strain exhibited significantly lower locomotor activity compared to the TU strain during both daytime and nighttime (Figure 4a-4c). Additionally, the AB strain displayed a significantly longer sleep duration than the TU strain (Figure 4d-4f). These findings highlight the distinct circadian rhythm behavior between two strains, with the AB strain showing reduced activity and extended sleep duration, underscoring the variability in circadian regulation.

**Figure 4.**
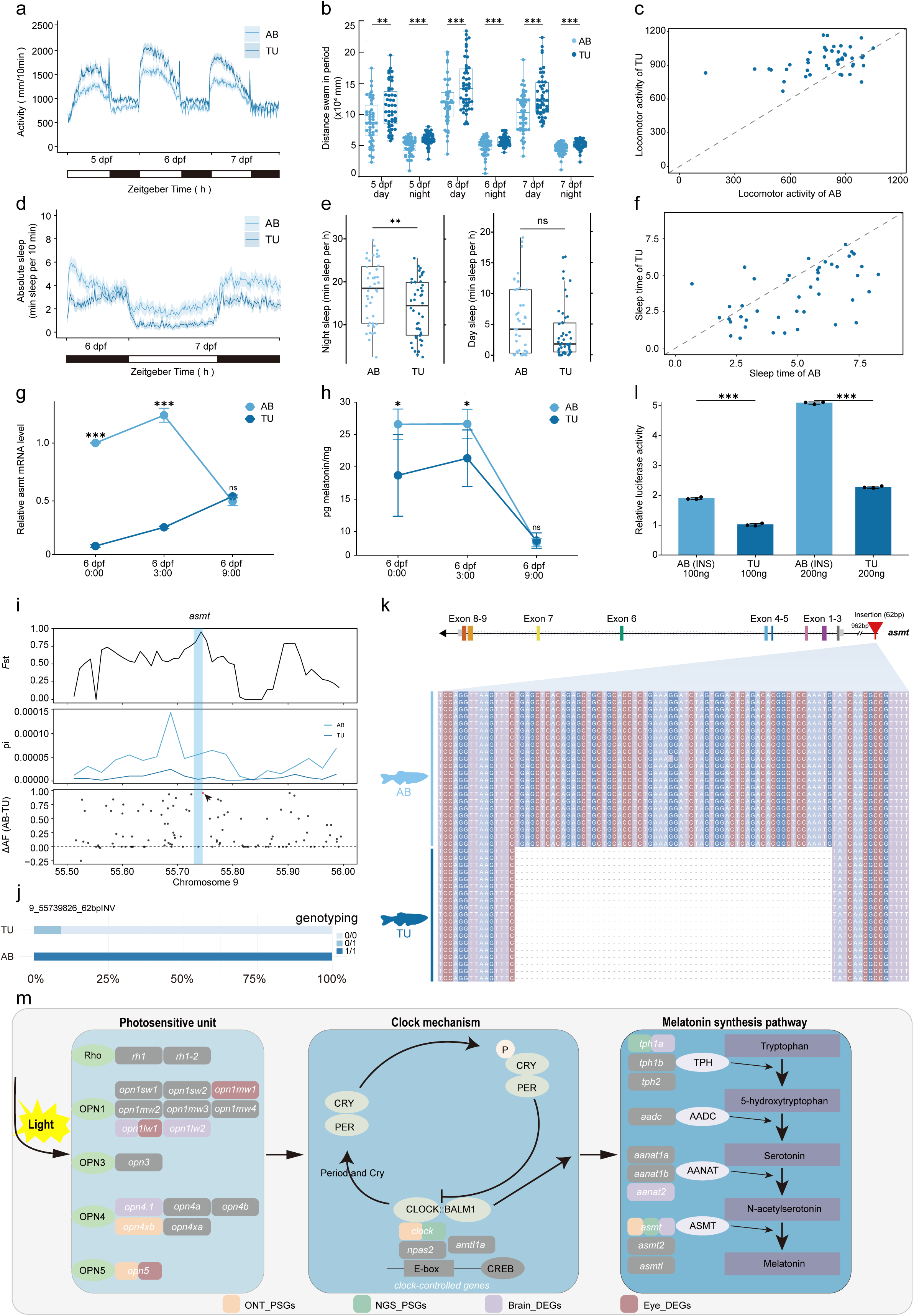
Effects of 62 bp insertion upstream of the *asmt* gene on the circadian rhythm phenotype among zebrafish strains. **a.** Locomotor activity nested per 10-min of AB strain and TU strain larvae from 5 dpf to 7 dpf under light and dark conditions. AB strain, n = 48; TU strain, n = 48. **b.** Quantification of locomotor activity in AB and TU strains at different developmental stages (5–7 dpf) during the day and night periods. **c.** Locomotor activity comparison between individual AB and TU larvae. Each point represents the locomotor activity of a single larva from the AB strain (x-axis) compared to the TU strain (y-axis). The dashed line represents the line of identity (y = x). **d.** Absolute sleep duration in AB and TU strains across the light/dark cycle at 6 and 7 dpf. The shaded regions represent the standard error of the mean. **e.** Box plots showing the spread of night and day sleep duration between AB and TU strains. **f.** Absolute sleep duration comparison between individual AB and TU larvae. Each point represents the locomotor activity of a single larva from the AB strain (x-axis) compared to the TU strain (y-axis). The dashed line represents the line of identity (y = x). **g.** qRT-PCR analysis of the mRNA level of *asmt* in AB and TU strains under light and dark condition. **h.** Comparison of melatonin content between AB and TU strains. **i.** Genomic analysis of the region around *asmt*. The top panel shows *F*st value, the middle panel shows π value, and the bottom panel shows ΔAF between AB and TU strains, highlighting a significant allele frequency shift at this locus (arrow), further suggesting selective pressure on the *asmt* region. **j.** The bar plot shows the distribution of genotypes. **k.** Sanger sequencing validation of the 62 bp insertion in the *asmt* gene. **l.** Luciferase signals and expression of two SV-haplotypes of *asmt*. **m.** Zebrafish circadian rhythm regulation mechanism and related essential genes. Genes highlighted in orange and green denote selections based on long-read and short-read data, respectively. DEGs in the brain and eye are marked in light purple and dark purple. The two-sided student’s t-test was applied and results were presented as mean ± SEM). *p < 0.05, **p < 0.01 and ***p < 0.001

To explore the genetic basis of these behavioral differences, we measured the expression levels of circadian rhythm-related genes. Given that *asmt* has been under selection across multiple methods and datasets, and it encodes the enzyme responsible for the final step of melatonin synthesis^51^, we performed quantitative real-time PCR (qRT-PCR) to assess *asmt* expression in AB and TU strain larvae (6 days post-fertilization [dpf]) at 0:00 a.m., 3:00 a.m., and 9:00 a.m. The result revealed that during the dark period (0:00 a.m. and 3:00 a.m.), *asmt* expression was significantly higher in the AB strain compared to the TU strain. However, the expression of *asmt* at the light period (9:00 a.m) showed no significant difference (Figure 4g). Concurrently, we measured melatonin content in larvae at the same time points. Consistent with the *asmt* expression data, melatonin levels were significantly higher in the AB strain during the dark period (0:00 a.m. and 3:00 a.m.), with no difference observed at 9:00 a.m. (Figure 4h).

Given the significant differences in *asmt* expression and melatonin levels, we focused on identifying genetic variants associated with circadian PSGs. Using both the SVs and SNPs datasets, we found that *asmt* resides in a region under strong positive selection, with prominent peaks in both *F*st and π near the gene (Figure 4i). The ΔAF (AB-TU) plot revealed a significant allele frequency shift at this locus. Among the identified variants, a 62-bp insertion was detected in the predicted promoter region, 962 bp upstream of *asmt*. Genotyping showed that the insertion was homozygous in 100% of AB individuals, while only 9.09% of TU individuals were heterozygosity for this variant (Figure 4j). Sanger sequencing validated the presence of this insertion, confirming its accuracy and ruling out sequencing artifacts (Figure 4k). Further analysis of publicly available chromosome-level genomes revealed that this variant was present exclusively in the AB strain (Supplementary Figure 6). To evaluate the functional impact of this insertion, we performed luciferase reporter assays, which demonstrated a significant increase in luciferase activity due to the 62-bp insertion (Figure 4l). These findings indicate that the insertion likely contributes to the observed strain-specific differences in *asmt* expression. To expand the scope of DEG screening, we collected brain and eye tissues from the two zebrafish strains at 3 a.m. and performed transcriptome sequencing. This analysis revealed several PSGs associated involved in circadian rhythm pathway that were differentially expressed between the strains (Figure 4m; Supplementary Table 12). These genes, represented in the circadian and melatonin synthesis pathways, form a complex regulatory network that may jointly contribute to the observed strain-specific differences in circadian rhythm phenotypes.

### Influence of positive selection on a non-synonymous mutation in the *mybbp1a* gene on zebrafish immunity

Many of the PSGs were enriched in immune-related pathways (Figure 5a; Supplementary Tables 8), leading us to hypothesize that immune capacity might differ between the two strains. To investigate this, we challenged AB and TU larvae 3 dpf with spring viremia of carp virus (SVCV) and photographed the larvae after 24 h. The dead larvae exhibited no movement, no blood circulation, and a degenerated body (Figure 5b, indicated by red arrows). We also counted the number of survivors at different time points and plotted survival curves (Figure 5c). These results showed that AB strain larvae are more resistant to SVCV infection compared to TU strain larvae (Figure 5b-5c). Furthermore, the typical antiviral gene *ifn* and pro-inflammatory cytokine genes, including *tnf*α, *il-1*β, and *il-6* were significantly enhanced in AB strain larvae compared to TU strain larvae (Figure 5d). Taken together, these findings indicate AB strain larvae have a stronger innate immune capacity against SVCV infection.

**Figure 5.**
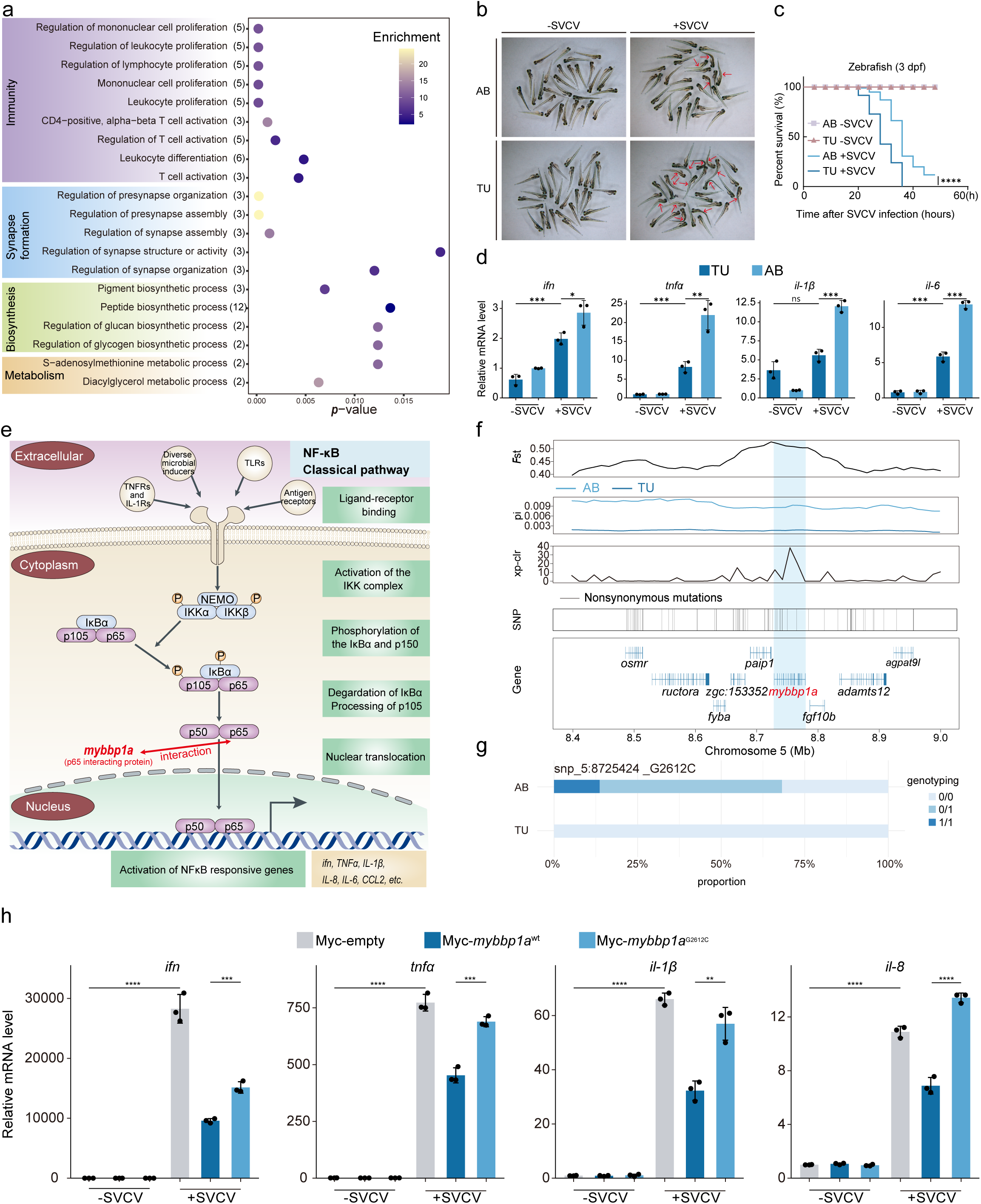
A non-synonymous mutation in the *mybbp1a* gene impairs innate immune response in zebrafish. **a.** Selected GO term enrichment for the genes under positive selection between two strains. The number of genes associated with each category is indicated in brackets after the term description, and enrichment values are indicated in colored scale. **b.** Representative images of AB and TU strain zebrafish larvae (3 dpf) post-SVCV infection. Mortality criteria include lack of movement, body curvature, and tissue degeneration (red arrows). **c.** Survival curve of zebrafish larvae (n = 96 per group) infected with or without SVCV for 44Lh. Statistical significance was assessed by log-rank test. **d.** qRT-PCR analysis of the mRNA levels of *ifn*, *tnf*α, *il-1*β, and *il-6* in TU strain and AB strain larvae after SVCV infection. **e.** Schematic diagram of the classic NF-kB pathway. **f.** The putative sweep region is additionally validated by *F*st, π, and xp-clr test on chromosome 5 (8.4 to 9.0 Mb). Gene annotations in the sweep region and SNPs nearly fixed for derived alleles in two strains are indicated at the bottom. **g.** Bar plot comparing genotype frequencies of the non-synonymous between two populations. **h.** qRT-PCR analysis of the mRNA levels of *ifn*, *tnf*α, *il-1*β, and *il-8* in EPC cells after SVCV infection. The two-sided student’s t-test was applied and results were presented as mean ± SEM). *p < 0.05, **p < 0.01, ***p < 0.001, and ****p < 0.0001.

The classical NF-κB pathway is known to play a key role in the immune response (Figure 5e)^52,53^. Among the PSGs, we identified *mybbp1a*, a gene whose encoded protein interacts with the p65 protein, a key subunit of the NF-κB complex^54,55^. This led us to hypothesize that genetic variations in *mybbp1a* may contribute to the immune differences between strains. Evidence supporting this hypothesis includes the large *F*st, XP-CLR, and π ratio values, along with numerous non-synonymous SNPs, all of which indicate that *mybbp1a* is under strong positive selection (Figure 5f). Notably, we discovered a non-synonymous mutation *mybbp1a*^G2612C^ exclusive to the AB strain (Figure 5g). To investigate whether *mybbp1a*^G2612C^ affects immune capacity, we overexpressed *mybbp1a*^wt^ and *mybbp1a*^G2612C^ in EPC cells followed by SVCV infection. We found that overexpressing *mybbp1a*^G2612C^ significantly upregulated SVCV induced expression of *ifn*, *tnf*α, *il-1*β, and *il-8* compared to overexpressing *mybbp1a*^wt^ (Figure 5h). These data supporting that the *mybbp1a*^G2612C^ mutation enhances immune response which likely contributes to the observed strain-specific immunity phenotypic differences.

## Discussion

This study provides the first comprehensive atlas of genetic variation in zebrafish model strains, achieved through an integrative analysis of population-scale genomic data. By combining SRS and LRS data from 22 AB and 22 TU zebrafish strains, we constructed a high-resolution variation landscape encompassing 39,580,820 SNPs, 10,614,893 InDels, and 178,894 SVs, expanding the reference for zebrafish genetic diversity research. Our analysis reveals three key findings: First, population structure analysis delineated two distinct genetic clusters corresponding to the AB and TU strains, reflecting significant phylogenetic divergence. Second, we explored the relationship between genetic variations and epigenetic regulation by integrating ATAC-seq and Hi-C data. Our results suggest that genetic variations may influence chromatin accessibility and TAD organization, potentially altering the local chromatin environment. Although these observations do not establish direct causality, they offer valuable insights into how genetic variations might affect epigenetic regulation. Third, genotype-phenotype association screening identified an SV and an SNP showing significant correlations with strain-specific traits, highlighting the functional consequences of these genetic variations. This work advances zebrafish genetics by 1) providing an essential repository of genetic variation for future studies, 2) revealing the evolutionary dynamics of multi-scale genomic variations, and 3) demonstrating how strain-specific genetic architectures influence phenotypic outcomes. Our results underscore the importance of considering genetic background differences when selecting zebrafish strain for biomedical research, particularly for studies aiming to establish genotype-phenotype correlations with high precision.

In the field of genetic architecture studies of zebrafish strains, multi-omics integration has made significant strides. Deng et al. performed de novo whole-genome assembly of individuals from the AB strain and systematically compared it with the GRCz11 reference genome (TU strain), successfully identifying several SVs associated with neurodevelopment and visual function^56^. However, their individual-level analysis limits the ability to explore the distribution patterns of genetic variation at the population level. Early SNP datasets from Victor Guryev et al. and Jaanus Suurväli et al., obtained using expressed sequence tags (EST) and restriction-site associated DNA sequencing (RAD-seq), respectively^12,13^, faced technical limitations in throughput—EST analysis identified only 50,000 SNPs, while RAD-seq detected 241,238 SNPs. These methods, therefore, fell short for a comprehensive analysis of inter-strain variation. Additionally, Kim H. Brown et al. and Yannick Schäfer et al. characterized CNVs using array comparative genomic hybridization (aCGH) and exome capture technologies, respectively^13,15^. However, aCGH has limited mapping resolution^57^, and exome capture designs fail to capture non-coding variations^58^. More importantly, existing studies have yet to establish a direct molecular link between genetic variations and phenotypic differences across laboratory strains.

By integrating SRS and LRS technologies, we systematically characterized the genome-wide variation landscape between zebrafish AB and TU strains. PCA and admixture analysis based on SVs dataset effectively distinguished AB from TU populations, aligning with differentiation patterns observed in SNPs dataset. Consistent with earlier studies reporting higher heterozygosity in AB^13^, our SNP-based analyses confirm this pattern and further show that AB also harbors higher π. In contrast, TU exhibits slower LD decay and a heavier ROH burden, with an excess of long (>1 Mb) tracts. This joint signature—low π, extended LD, and long ROH—is more parsimoniously explained by a smaller long-term effective population size and more recent inbreeding in TU, plausibly reflecting colony history. These demographic differences provide a baseline for all downstream scans and help explain why signal breadth and mapping resolution differ between strains.

Three-dimensional (3D) genome architecture is crucial for regulating gene expression, as it facilitates the spatial arrangement of distal cis-regulatory elements (CREs) in proximity to their target genes ^59–61^. TADs are large genomic regions where sequences within the domain interact more frequently with sequences outside the domain^62–64^. These domains are separated by TAD boundaries, which act as barriers to prevent interactions between CREs and promoters located in different TADs ^65^. The organization of these domains is thought to play a pivotal role in regulating gene expression by controlling the accessibility and interaction of CREs with their target promoters. Studies have shown that SNPs may regulate CRE activity, thereby influencing the transcription of target genes ^66,67^, while SVs may lead to 3D genome rearrangements that alter regulatory relationships, leading to subsequent changes in gene expression ^68–70^. By integrating our variation dataset with epigenetic data, we found that SNPs associated with strain-specific differentiation were enriched in dACRs, suggesting these SNPs may modulate CRE activity and contribute to strain-specific differentiation. Furthermore, we found that SVs were more likely to occur in strain-specific TADs, accompanied by a decrease in insulation scores, indicating that these SVs may perturb TAD structures and disrupt regulatory networks, thereby affecting gene expression. These results support the model that variation between strains is closely related to epigenetic inheritance. Specifically, (1) dACR regions are more likely to be associated with highly differentiated SNPs; (2) SVs tend to perturb TAD structures, reshaping regulatory relationships and influencing gene expression. Notably, we detected significant spatial concordance of selection signals among SNPs, InDels and SVs. By examining genes associated with SNPs, InDels, and SVs exhibiting prominent *F*st values in both populations, we identified several genes that were linked to all three types of variations, suggesting that these genes may play a role in strain differentiation. In contrast, other genes were linked to only one or two types of variation, indicating that different types of genetic variation may contribute synergistically or in parallel to driving the differences between the strains^71^.

Building on the genetic variation data between the strains, we further explored the potential associations between these variations and phenotypic differences. The hierarchical regulatory system of the circadian rhythm consists of three core components: the environmental time signal sensing module, the core molecular oscillator regulatory network, and the downstream physiological output pathways^72–74^. Our study identifies significant positive selection signals in melanopsin (*opn4xb*) and neuropsin (*opn5*) across strains, with *opn5* exhibiting strain-specific differential expression patterns in the eye. Previous studies gene knockout experiments have demonstrated that both of these genes play a critical regulatory role in photoperiod adaptation^75–78^. Among the components of the core molecular oscillator, only *clock1a*, which encodes a key regulatory element of the core transcription-translation negative feedback loop^79^, is under positive selection. Notably, within the melatonin biosynthesis pathway, the tryptophan hydroxylase gene *tph1a* is not only under positive selection but also shows strain-specific differential expression in brain tissue^80^. Additionally, the rate-limiting enzyme gene *aanat2* exhibits significant strain-specific expression differences in brain^81^. The terminal enzyme gene *asmt*, also under strong positive selection, shows differential expression patterns in the brain across strains. Our analysis identified a 62-bp strain-specific insertion upstream of *asmt*. qRT-PCR analysis demonstrated the mRNA level of *asmt* was significantly upregulated in the AB strain, correlating with circadian-related phenotypic divergence: the AB strain exhibited enhanced melatonin oscillation amplitude, locomotor activity rhythm, and longer sleep duration. Dual-luciferase assays confirmed the enhancer function of this insertion. Thus, a possible mechanistic explanation is that the strain-specific 62-bp insertion in AB enhances *asmt* expression, leading to increases melatonin production, which in turn contributes to reduced locomotor activity rhythm and longer sleep duration. However, definitive conclusions on causality would require targeted gene editing. Through this analysis, we identified several key genes across all three core components of the circadian rhythm network. These genes may collectively drive the observed differences in the circadian rhythm phenotype between the strains.

SVCV infection experiments reveal significant differences in the efficacy of the innate immune response between the AB and TU zebrafish strain. Positive selection analysis identified strong signatures at the *mybbp1a* locus, which encodes an interacting partner of the NF-κB p65 subunit. The strain-specific c.2612G>C (p.Gly871Ala) missense mutation showed fixation in AB strain. In vitro functional characterization demonstrated that this mutation significantly increased proinflammatory cytokine secretion upon SVCV stimulation, suggesting its role in mediating inter-strain infection resistance through modulation of NF-κB signaling dynamics. Notably, among the positively selected genes, synaptic plasticity regulatory genes such as *glra2*, *cpeb3* and *gng8* show significant differentiation between the strains. Experimental data from multispecies knockout models reveal critical roles of *gng8*, *glra2*, and *cpeb3* in neurocognitive functions. For instance, *gng8*-deficient murine models exhibit significant impairments in spatial learning and memory retention^82^. Similarly, *glra2* ablation induced conserved phenotypic manifestations across zebrafish and murine models, including abnormal axonal arborization and compromised working memory performance^83^. Additionally, *cpeb3* knockout mice show deficits in hippocampal-dependent memory consolidation, particularly in contextual fear conditioning tests^84^. Building on previous studies of inter-strain differences in learning and memory abilities in zebrafish^18^, we hypothesize that the genetic variations associated with these genes may contribute to phenotypic differences in learning and memory.

This study represents the first high-resolution, whole-genome variation map of the AB and TU strain populations of zebrafish, filling a significant gap in the field and providing new insights into the relationship between genetic and phenotypic variation. We emphasize the importance of considering strain-specific genetic backgrounds in experimental design, particularly in studies of circadian rhythms, immune response, and cognitive functions, as these factors can significantly influence phenotypic outcomes.

## Methods

### Sample collection for Nanopore LRS and Illumina SRS and DNA extraction

A total of 44 unrelated, healthy zebrafish (*Danio rerio*), with an average age of 6 months, were obtained from the China Zebrafish Resource Center. The group included 11 individuals from each of the following categories: male and female AB (ZFIN ID: ZDB-GENO-960809–7) strains, and male and female Tübingen (TU; ZFIN ID: ZDB-GENO-990623–3) strains. High-quality genomic DNA was extracted from muscle tissue after skin removal using an SDS-based method. RNA contamination was removed using RNase A. DNA quality was assessed by agarose gel electrophoresis, and the integrity of the DNA was verified by visualization of intact DNA bands.

### Library construction and sequencing of whole genome-seq

For Illumina sequencing, DNA libraries (350 bp) were constructed for each sample according to the manufacturer’s instructions. Sequencing was subsequently performed on an Illumina NovaSeq platform with 150-bp paired-end reads, achieving an average sequencing depth of 31×.

For Nanopore sequencing, DNA repair, end repair, and adapter ligation were performed during library preparation. A total of 2 µg of DNA was fragmented using a g-TUBR (Covaris). DNA repair was carried out using the NEBNext FFPE DNA Repair Mix (M6630L, NEB), followed by end repair with the NEBNext Ultra II End Repair/dA-Tailing Module (E7546L, NEB). Adapter ligation was conducted using the NEBNext Blunt/TA Ligase Master Mix (M0367L, NEB) and the Ligation Sequencing Kit 1D (SQK-LSK109, Oxford Nanopore Technologies). DNA concentration was quantified using a Qubit Fluorometer 2.0 (ThermoFisher Scientific, Waltham, MA). Long-read sequencing was performed on a PromethION sequencer with a 1D flow cell and protein pore R9.4.1 1D chemistry, in accordance with the manufacturer’s guidelines. Base-calling was conducted in batches using guppy (v3.2.8) with default parameters. Each genome was sequenced to a coverage depth ranging from 19× to 33×, with an average depth of 24×.

### Detection and annotation of SNP and InDel

After quality control of the SRS data, clean reads were aligned to the *Danio rerio* reference genome (GRCz11) using BWA-MEM (v0.7.19)^85^ with default parameters. The resulting alignment files were then sorted using SAMtools (v1.16)^86^, and duplicate reads were removed with Picard Tools (v3.3.0) (https://github.com/broadinstitute/picard). SNPs and InDels were identified using the Genome Analysis Toolkit (GATK, v4.4.0.0-0)^87^, employing the “HaplotypeCaller”, “CombineGVCFs”, “GenotypeGVCFs”, and “SelectVariants” tools. After variant calling, the data were filtered using “SelectVariants” with the following criteria: “QUAL < 30 || QD < 2.0 || MQ < 40.0 || FS > 60.0 || SOR > 4.0 || MQRankSum < -12.5 || ReadPosRankSum < -8.0”. Finally, SNPs and InDels were annotated using ANNOVAR^88^.

### Detection and genotyping of SV

Raw long reads were aligned to the Danio rerio reference genome (GRCz11) using minimap2 (v2.26-r1175)^89^ with the parameters “-Y --MD -t 16 -ax map-ont”. SAMtools (v1.16)^86^ was used to sort and convert alignments into BAM format. To generate a high-quality set of SVs, two SV calling programs, CuteSV (v2.0.2)^90^ and Sniffles2 (v2.0.7)^91^, were employed. SVs were called using Sniffles with the pareameters “-t 5 --minsvlen 50 --minsupport auto --mapq 20” and cuteSV with the parameters “-t 10 -S hg002 --min_size 50 --max_size 100000 --retain_work_dir --report_readid –genotype --max_cluster_bias_INS 100 --diff_ratio_merging_INS 0.3 --max_cluster_bias_DEL 100 --diff_ratio_merging_DEL 0.3”). The SV callsets from both callers were merged for each SV type using SURVIVOR (v1.0.7)^92^ with the parameters “merge 1000 2 1 1 0 50”. Subsequently, CuteSV was re-run across all samples with the parameters “-t 10 -S sample --min_size 50 --max_size 100000 --retain_work_dir --report_readid --min_support 3 --genotype -Ivcf merge_vcf --max_cluster_bias_INS 100 --diff_ratio_merging_INS 0.3 --max_cluster_bias_DEL 100 --diff_ratio_merging_DEL 0.3”, and the resulting SVs were again combined using SURVIVOR.

SVs identified from LRS data were genotyped using SRS data generated in this study. All short reads were mapped to GRCz11 zebrafish reference by BWA-MEM with default parameters. Using SVs from LRS data, we employed Paragraph (v 2.4a)^93^ to genotype the combined SVs from SRS data and all genotypes which failed to pass filter by Paragraph were placed with missing genotypes (./.). Genotyped SVs were subsequently filtered using Plink2 (v2)^94^ with the parameters “--vcf vcf --allow-no-sex --maf 0.05 --geno 0.2 --r2-phased allow-ambiguous-allele --ld-window 300 --ld-window-r2 0 --autosome-num 25 --threads 8”. Finally, the Variant Effect Predictor (VEP, v103)^95^ was used to annotate the SVs.

### Population structure and genetic diversity analysis

Population structure is constructed based on SV datasets. A phylogenetic tree was constructed using the neighbor-joining method with PHYLIPNEW (v3.69.650)^96^, and the unrooted tree was visualized with iTOL (https://itol.embl.de/). Principal component analysis (PCA) was performed using GCTA (v1.93)^97^, and population structure analysis was carried out using ADMIXTURE (v1.3.0)^98^.Genetic diversity was assessed using SNP datasets. Nucleotide diversity (π) was calculated with vcftools (v0.1.13)^99^ using the following parameters: “vcftools --vcf snp.vcf --fst-window-size 100000 --fst-window-step 10000 --keep pop1 --out π_pop1”. Individual heterozygosity was estimated using PLINK and vcftools. Tajima’s D value was also calculated using vcftools.

### Linkage disequilibrium (LD) and estimation of inbreeding

To assess the degree of linkage disequilibrium (LD) between the AB and TU strains, PopLDdecay (v3.42)^100^ was used to calculate the coefficient of determination (r^2^) and generate LD decay curves.Inbreeding depression was tested by calculating runs of homozygosity (ROH) using PLINK with the following parameters: “--homozyg-window-snp 50 --homozyg-snp 50 --homozyg-kb 500 --homozyg-density 50 --homozyg-gap 1000 --homozyg-window-missing 5 --homozyg-window-threshold 0.05 --homozyg-window-het 3”. The individual inbreeding coefficient, *F*_ROH,_ was calculated as the ratio of the ROH segment to the total length of the 25 autosomes.

### Library construction and sequencing of ATAC-seq

The 6 dpf zebrafish were first washed with a 0.09% NaCl solution and then homogenized into a fine powder in liquid nitrogen. Lysis buffer was added to the powdered zebrafish tissue and incubated for 10 minutes at 4°C on a rotating mixer. The cell suspension was filtered through a 40 µm cell strainer and washed three times with cold PBS buffer. ATAC-seq assays were performed by Wuhan GeneRead Biotechnology Co., Ltd. After assessing the purity and integrity of the nuclei under a microscope, approximately 50,000 nuclei were used for tagmentation according to standard protocols. Tn5-transposed DNA was purified using AMPure DNA magnetic beads. A PCR reaction was conducted on a portion of the DNA to determine the optimal number of PCR cycles (average 11). The quality of the amplified libraries was assessed using an Agilent TapeStation 2100 with a D5000 DNA ScreenTape to visualize nucleosomal laddering. Biological replicates were performed in duplicate for all ATAC experiments. The final library was sequenced on the Illumina NovaSeq XPlus platform (San Diego, CA, USA) in PE150 mode.

### ATAC–seq data analysis and dACR identification

Raw ATAC-seq data were processed using a custom pipeline. BAM files were mapped to GRCz11, and Tn5 insertion sites were identified using MACS2 (v2.2.9.1)^101^. High-frequency peaks were generated for each sample and filtered by chromosome range (chr1–25) using bedtools (v2.28.0-33-g0f45761)^102^ intersect. A fixed width of 500 bp around the summit was applied to generate fixed-width peaks. To create a blacklist of high-frequency and extreme signal peaks, two steps were employed: first, high-frequency peaks, defined as those appearing in ≥80% of the samples, were identified using MACS2 and bedtools merge, and these peaks were intersected with allowed chromosomes. Second, extreme high-signal peaks were identified by normalizing read counts to CPM using bamCoverage from deepTools (v3.5.6)^103^, with peaks exceeding the 99.9th percentile of signal intensity selected as high-signal peaks. The final blacklist was generated by intersecting the high-frequency and high-signal peak sets using bedtools intersect, followed by merging overlapping regions. Peaks were called for each sample using MACS2, and the resulting raw peaks were processed into clean peaks by removing regions that overlapped with the blacklist. The clean peaks were then used for downstream differential analysis and ACR identification. dACRs were identified using the eFDR filtering method according to previous study^104^. First, Tn5 insertion sites were extracted from the mapped data, and the coverage of Tn5 sites in the fixed500.clean.bed regions was calculated using bedtools coverage. Random background regions were generated with bedtools shuffle, which excluded ACRs from the selection space. The true peaks and random peaks were processed using the eFDR.R script to remove regions with low accessibility.

### HiC data analysis

The Hi-C raw data were downloaded from PRJNA734711 (AB strain)^56^ and PRJNA553572 (TU strain)^105^.Low-quality reads were filtered out using Trimmomatic (v0.39)^106^. Paired-end reads were then aligned to the GRCz11 genome using Bowtie2 (v 2.5.2)^107^. After alignment, singleton, multi-mapped, dumped, dangling, self-circle paired-end reads, and PCR duplicates were removed via HiC-Pro (v3.1.0)^108^. Contact matrices were constructed with varying bin sizes (10 kb, 25 kb, 40 kb, 100 kb). The raw contact maps were subsequently normalized using an iterative correction method based on a sparse matrix implementation. TADs and boundaries were identified at a resolution of 40 kb using hicFindTADs in the HiCExplorer suite (v3.7.5)^109^. Hi-C loops were detected from the matrices using HiCCUPS^110^ with parameters: “-ignore-sparsity -r 5000, 10000, 25000 -k KR,”. The Hi-C matrices were visualized using the pyGenomeTracks tool^111^.

### Detection of population selective signals

To investigate positively selected regions in the AB and TU strains, three methods were employed: the cross-population composite likelihood ratio test (XP-CLR), π ratio and *F*st statistics. Both SNPs and InDels were analyzed separately using these methods. *F*st values were calculated for both SNP and InDel datasets using vcftools with the following parameters: “vcftools --vcf snp.vcf --fst-window-size 100000 --fst-window-step 10000 --weir-fst-pop pop1 --weir-fst-pop pop2 --out pop1_pop2.fst”. The π ratio was computed as the ratio of π (AB) to π (TU) and π (TU) to π (AB) for each corresponding window. XP-CLR values were calculated using xpclr (v1.1.2)^112^ with a window size of 100,000 and a window step of 10,000. The top 5% of ranked windows based on *F*st, π ratio and XP-CLR scores were considered candidate regions under selection. The intersection of these selected regions with the General Feature Format 3 file was obtained using bedtools, and the genes within these intersecting regions were defined as positively selected genes (PSGs). To identify population-stratified SVs, *F*st and the π ratio were calculated for SVs between the AB and TU strains. Genomic windows within the top 5% of both *F*st and π ratio were defined as candidate regions under selection. Using a similar approach to that for SNP analysis, selected genes associated with SVs were identified by intersecting SV regions with gene annotations in the GFF3 file using bedtools. Additionally, genes located within 5 kb upstream and downstream of each SV were also considered potential targets of selection. The overlap between these two gene sets was used to determine positively selected genes associated with SVs.

### Behavioral assays

Zebrafish larval behavior was monitored following established protocols with slight modifications. Embryos were maintained at 28°C under a 14-hour light and 10-hour dark (LD) cycle during the first ∼96 hours of development. Larval locomotor activities were recorded from days 5 to 7 post-fertilization using an automated video tracking system (Zebrabox, Viewpoint LifeSciences) and analyzed with Zebralab software (v 2.30).

On the night of 4 dpf, individual larvae were transferred into the wells of a 24-well plate, which was then placed inside the Zebrabox for acclimation. A 24-well plate was chosen for its ability to allow greater larval activity compared to 96-well plates. The Zebrabox was configured to provide continuous infrared illumination, with white light exposure from 8:00 a.m. to 10:00 p.m. under LD conditions. The total distance moved by each larva during each period was recorded using Tracking mode, which measured the larvae’s movement throughout the experiment. Each experiment included 12 AB and 12 TU larvae, and the process was repeated four times for reproducibility. Sleep duration measurement was performed by recording larvae in quantification mode with the following settings: detection sensitivity set to 25, burst set to 20, freeze set to 5, scale set to 20, and the transparent checkbox selected. The integration period was set to 60 seconds, and sleep periods were defined as 1 minute of immobility during the recording^113^. The data were integrated using a custom Perl script.

### Variations Validated by PCR and Sanger Sequencing

To ensure the reliability of the detected variations, we employed polymerase chain reaction (PCR) followed by Sanger sequencing for verification. Genomic DNA was extracted from the muscle tissue of zebrafish strains AB and TU. PCR reactions were performed using 2× Taq Master Mix (VAZYME). The insertion in the *asmt* gene was confirmed through PCR genotyping with the following primer pairs: (F: GTTCTTCAAACTGGGCTCG, R: ACCCTCGTTGTGGCTGTC).

### Quantitative real-time PCR

Total RNA from the larvae and cells were extracted using TransZol Up (TransGen Biotech) following the manufacturer’s instructions. Complementary DNA (cDNA) was synthesized using the RevertAid First Strand cDNA Synthesis Kit (Thermo Scientific). qRT-PCR assays were performed using the MonAmp SYBR Green qPCR Mix (High Rox) (Monad Bio). The primers used for qRT-PCR assays are provided in the supplementary table 14.

### ELISA

The concentration of melatonin (MT) in the serum was measured using a commercially available ELISA kit (Mlbio, Cat. No. ml025846-1) according to the manufacturer’s instructions. The melatonin concentration was determined based on the standard curve.

### RNA-seq analysis

Based on the q RT-PCR results of rhythmic gene expression, we selected 3:00 a.m. as the sampling time to collect the brains and eyes from two zebrafish strains for transcriptome sequencing. Total RNA was extracted using TRIzol reagent (TIANGEN) following the instructions. The concentration of extracted nucleic acid was detected with Nanodrop2000, and the integrity was detected with Agient2100. We used 1-2 μg RNA per sample to construct the cDNA library and using Qubit 3.0 perform preliminary quantification. The library was then sequenced using Illumina NovaSeq 6000 and 150 bp paired-end reads were produced. Raw reads were processed with fastp (v 0.23.4)^114^ to remove low-quality reads. The clean reads were then aligned to the GRCz11 reference genome using HISAT2 (v 2.1.0)^115^, and transcripts were assembled and quantified using StringTie (v 2.2.2)^116^. Differential expression analysis was conducted using the DESeq2 R package (v 1.36.0)^117^. Genes with a p-value < 0.05 and |Log2FC| > 1.25 were identified as differentially expressed between AB and TU strain.

### Plasmid constructions

Zebrafish *mybbp1a* (Gene ID: ENSDARG00000078214), the *mybbp1a*^G2612C^ mutant (p.Gly871Ala), and the *asmt* promoter with or without the SV region fragments (Supplementary Table 15) were amplified by PCR. The amplified genes were then cloned into the pCMV-Myc vector. The *asmt* promoters, both with and without the SV region, were cloned into the PGL3-basic vector. All constructs were confirmed by Sanger sequencing. The plasmids were transfected into cells using VigoFect (#T001, Vigorous Biotech, Beijing, China).

### Cells and virus

Zebrafish liver (ZFL) cells (originally from the American Type Culture Collection) were cultured in Ham’s F-12 medium (Gibco) supplemented with 10% fetal bovine serum (FBS) (VivaCell) at 28°C in a humidified incubator containing 5% CO_2_. Epithelioma papulosum cyprini (EPC) cells (originally from the American Type Culture Collection) were maintained in medium 199 (VivaCell) containing 10% FBS at 28°C in a humidified incubator containing 5% CO_2_. Spring viremia of carp virus (SVCV) was propagated in EPC cells until the cytopathic effect (CPE) was complete, and the culture medium was collected and stored at -80°C until use.

### Luciferase reporter assays

ZFL cells were cultured in 24-well plates and transfected with varying amounts of plasmids using VigoFect (Vigorous Biotech, Beijing, China). CMV-Renilla was co-transfected as an internal control. Following the designated transfection period, luciferase activity was quantified using the Dual-Luciferase Reporter Assay System (Promega). Data were normalized to Renilla luciferase.

### Viral infection

For viral infection of the AB and TU strain larvae, 30 larvae each group were placed in 60 mm disposable cell culture dishes containing 4 mL of egg water and 1 mL SVCV (approximately 6 × 10^7^ TCID_50_/mL). For the survival curve assay, zebrafish larvae were placed in 96-well plates. One larva per well was individually infected with SVCV. After infection, the remaining survival larvae in each group were monitored at indicated time. For viral infection of EPC cells, indicated plasmids were transfected to EPC cells for 24 h following infected with SVCV for 16 h.

### Statistical analysis

Results with error bars represent mean ± SEM. Two-sided Student’s t-test was used for statistical analysis. Survival analysis was performed using the log-rank test to assess statistical significance. Statistical significance is denoted as follows: *p < 0.05, **p < 0.01, ***p < 0.001, and ****p < 0.0001.

## Acknowledgements

This work was supported by the National Natural Science Foundation of China (32422010), Youth Innovation Promotion Association, Chinese Academy of Sciences (http://www.yicas.cn), and the Young Top-notch Talent Cultivation Program of Hubei Province to L.Y. We would like to thank Dr. Zhixian Qiao and Xiaocui Chai at The Analysis and Testing Center of Institute of Hydrobiology, Chinese Academy of Sciences for their guidance on the zebrafish behavioral experiments.

## Author contributions

S.H. lead the study. L.Y. designed the research. W.C., J.J., Y.L., and L.Z. performed data analyses. J.W. and Z.L performed functional experiments. W.C. and L.Y. wrote the manuscript.

## Ethics Statement

All experiments performed in this study were approved by the Institutional Animal care and Use Committee of Institute of Hydrobiology, Chinese Academy of Sciences (IHB, CAS, Protocol NO. 2016-018).

## Data availability

All the raw sequencing data have been deposited at the NCBI under BioProject number PRJNA1232966.

